# Structural basis of ethnic-specific variants of PAX4 associated with type 2 diabetes

**DOI:** 10.1101/2020.03.24.006254

**Authors:** Jun Hosoe, Ken Suzuki, Takashi Kato, Yukinori Okada, Momoko Horikoshi, Nobuhiro Shojima, Toshimasa Yamauchi, Takashi Kadowaki

## Abstract

Recently, we conducted genome-wide association studies of type 2 diabetes (T2D) in a Japanese population, which identified 20 novel T2D loci that were not associated with T2D in Europeans. Moreover, nine novel missense risk variants, such as those of *PAX4*, were not rare in the Japanese population, but rare in Europeans. We report *in silico* structural analysis of ethnic-specific variants of *PAX4*, which suggests the pathogenic effect of these variants.

## Main Text

Large-scale meta-analyses of genome-wide association studies (GWAS) of type 2 diabetes (T2D) have been performed recently^1-4^. In particular, Mahajan et al. reported T2D GWAS meta-analysis data on approximately 900,000 individuals of European ancestry, increasing the association signals in over 240 loci associated with T2D^3^. Recently, we conducted an extensive T2D GWAS meta-analysis in a Japanese population and identified 28 new loci^4^, highlighting the value of genetic research in ethnically diverse populations. As an example of clinical differences between ethnic populations, East Asians are more likely to develop T2D at a lower body mass index than Europeans^5^, who have been reported to show a higher insulin response and lower insulin sensitivity.

We demonstrated that the biological pathway of the maturity onset diabetes of the young (MODY), monogenic diabetes characterized by reduced β-cell function, showed the most significant association with T2D in both Japanese and European populations^4^, by performing pathway analysis of GWAS summary data in the two populations^1,4^ using Pascal software. Here we focused on protein-coding genes involved in the MODY pathway mapping nearest to lead variants at T2D loci, as previously reported^6^, and investigated ethnic differences in the associations of these loci with T2D between Japanese and European populations^3,4^. As suggested by a previous study^4^, most of these lead variants at T2D loci in either the Japanese or European population showed at least nominally significant (*P* < 0.05) associations in the alternative population, even if not genome-wide significant, except for variants that were rare or monomorphic in one population such as *NKX6-1* rs201597274, *HNF1A* rs187150787, and *PAX4* rs2233580 in the Japanese population and *HNF1A* rs56348580 in the European population. Interestingly, there were several loci such as *GCK* where lead variants in these populations were independent of each other (**Supplementary Table 1** and **2**). As an example of a locus specific to East Asians, a previously unreported missense T2D variant of *PAX4* NM_006193:c.574C>A:p.(Arg192Ser) reached genome-wide significance^4^. This variant was located at the same amino acid as another established independent T2D variant NM_006193:c.575G>A:p.(Arg192His) (Table 2 in ref. ^4^). *PAX4* encodes a transcription factor that is important for β-cell development, and rare mutations of this gene are suggested to cause a subtype of MODY (OMIM #612225). Arg192 of PAX4 is located in the homeodomain, which is a DNA-binding domain conserved in a large family of transcription factors. We conducted an *in silico* structural analysis of the pathogenic effect of these *PAX4* variants, which revealed a decrease of the PAX4 homeodomain stability and reduced DNA binding by this domain in both variants (**Fig. 1** and **2, Supplementary Note**).

**Fig. 1.**
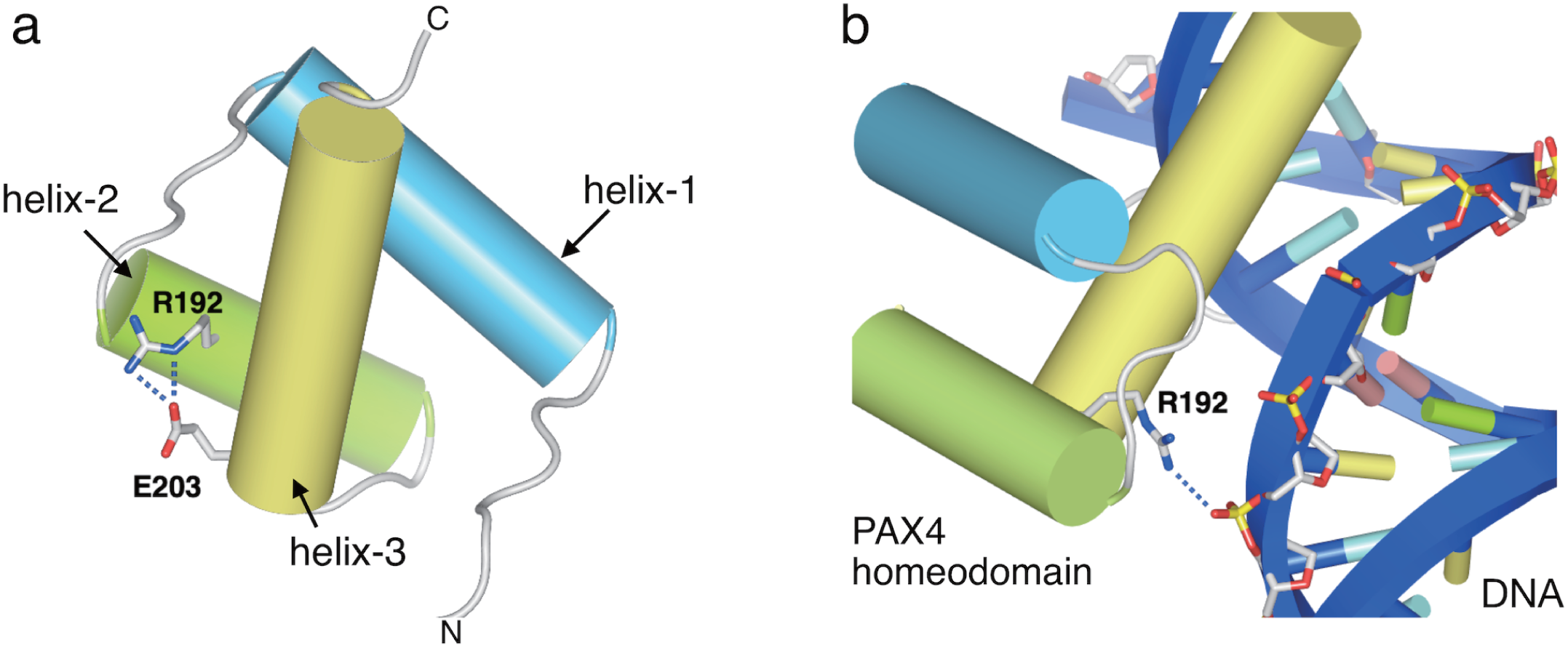
Structural model of the homeodomain in human PAX4. **a** The human PAX4 homeodomain is represented as cylinders, with the Arg192 and Glu203 residues being shown as sticks. The PAX4 homeodomain consists of three helices: helix-1 (light-blue), helix-2 (green), and helix-3 (yellow). Arg192 located on helix-2 is predicted to form salt-bridges with Glu203 located on helix-3. **b** Binding of the human PAX4 homeodomain to double-stranded DNA. DNA is shown by ribbon representation in blue. Arg192 of PAX4 (sticks) is predicted to bind with the phosphate backbone (sticks) of DNA.

**Fig. 2.**
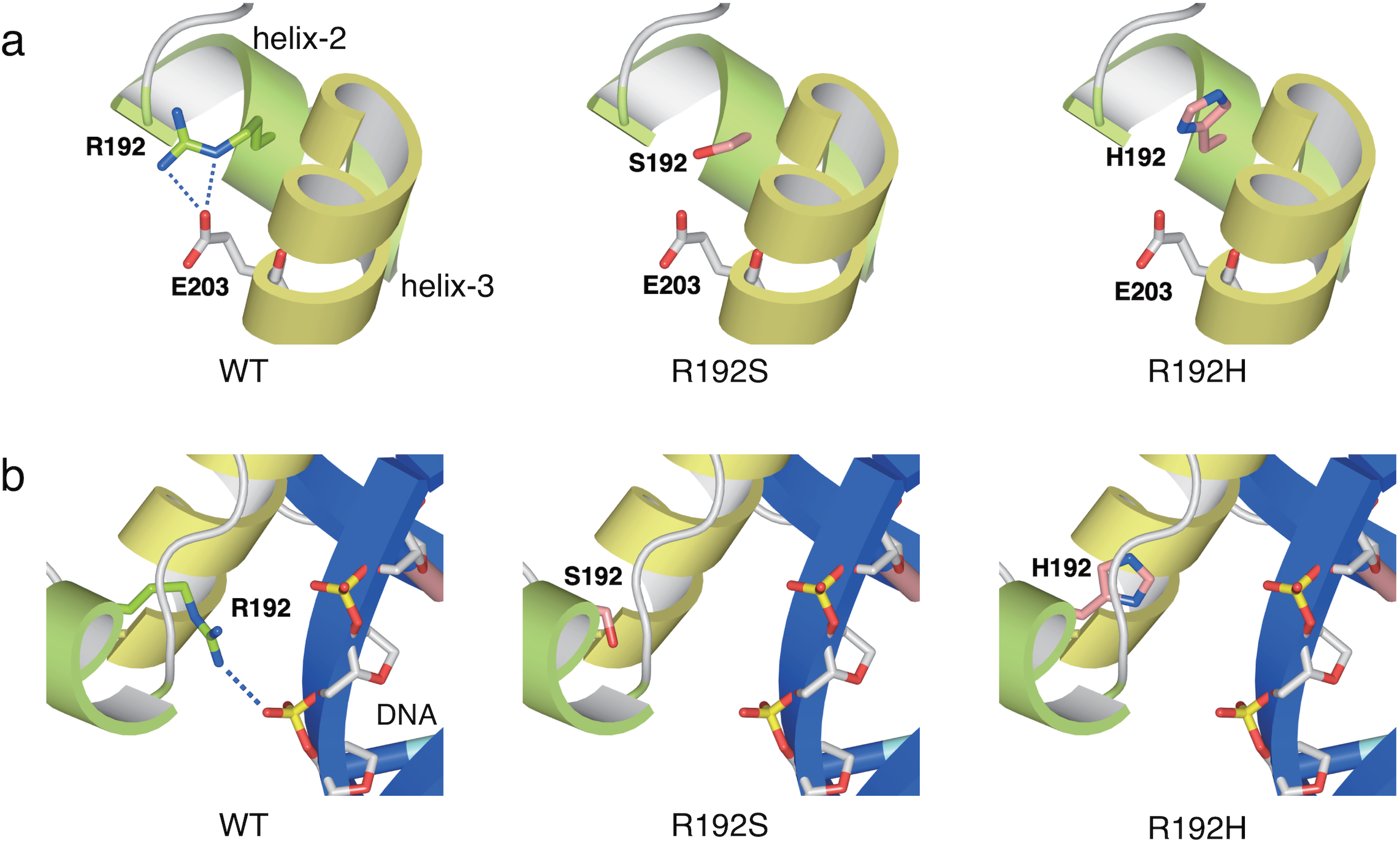
Structural analysis of two PAX4 variants, p.Arg192Ser and p.Arg192His. PAX4 are shown by ribbon representation, with the key residues of PAX4 as sticks. The helix-2 and the helix-3 of PAX4 are shown in green and yellow, respectively. Wild-type Arg192 is green, while the Ser192 and His192 mutants are light red. **a** In wild-type PAX4, Arg192 forms salt-bridges with Glu203 (dotted blue line). In both mutants (p.Arg192Ser and p.Arg192His), formation of salt-bridges with Glu203 is disrupted. **b** DNA is shown by ribbon representation in blue, with the phosphate backbone as sticks. In wild-type PAX4, Arg192 directly binds to the phosphate backbone of DNA (dotted blue line). In both mutants (p.Arg192Ser and p.Arg192His), binding with the phosphate backbone of DNA is disrupted.

To physiologically classify T2D loci, cluster analyses have been performed using data on metabolic traits^1,2,7^. Mahajan et al. performed a hierarchical cluster analysis of loci associated with T2D in ethnically diverse populations, generating three main clusters associated with BMI/dyslipidemia, insulin secretion, and insulin action^2^. The cluster related to reduced insulin secretion contained a number of important loci involved in β-cell function as shown in their Supplementary Fig. 6^2^, with several lead variants, such as those of *PAX4* (p.Arg192His) and *HNF1A*, that showed distinct minor allele frequency (MAF) spectra and ethnically different associations (Supplementary Table 10 and 11 in ref. ^2^). Such differences could lead to distinct proportions of the clusters of T2D loci with different underlying biological pathways^2,7^, which may result in variable proportions of subgroups of T2D patients classified based on their genetics in ethnically diverse populations. For example, the *PAX4* variants are specific to East Asians^4^, which presumably increase the proportion of the cluster of T2D loci with biological pathways influencing β-cell function in East Asian populations.

In a recent review article^8^, the pronounced ethnic heterogeneity associated with ethnic-specific risk variants (e.g., *PAX4* p.Arg192His and *HNF1A* p.Glu508Lys^9^) was suggested to be relatively unusual and not to explain observed ethnic differences in presentation of T2D. It was also suggested that rare ethnic-specific variants identified through sequencing studies may be more important in this regard^8^. Whereas T2D GWAS meta-analysis recently performed in a Japanese population identified no less than 20 novel T2D loci^4^ that were not associated with T2D in Europeans^1,3^ (Supplementary Table 3 in ref. ^4^). Ethnic differences in the associations of a number of new loci with T2D may be affected by several factors, such as differences in allele frequency and diverse patterns of linkage disequilibrium in these populations as suggested recently^10^. For example, we identified nine novel missense variants such as those of *PAX4* (p.Arg192Ser) and *GLP1R* (p.Arg131Gln)^4^ that were in linkage disequilibrium with the lead variants of T2D loci and not rare in Japanese population, but rare or monomorphic in Europeans (Supplementary Table 6 in ref. ^4^).

Considering the differences in sample size between these studies^3,4^, more large-scale genetic studies of East Asian populations are needed to fully elucidate similarities and differences in the pathogenesis of T2D between them. In conclusion, our new GWAS data^4^ suggested that large-scale single ethnic genetic studies could be useful for identifying ethnic-specific risk variants including relatively common ones, providing new insights into the genetic heterogeneity of T2D in diverse ethnic groups. The new information thus obtained would facilitate development of tailored therapy for this disease in various populations.

## Acknowledgements

We acknowledge all authors of our GWAS meta-analysis of T2D in a Japanese population^4^ for their contribution to advancing genetic research. This study was supported by a grant-in-aid for scientific research from the Ministry of Education, Culture, Sports, Science and Technology of Japan (MEXT) (grant 19K16534 to J.H.). This research was also supported by Advanced Genome Research and Bioinformatics Study to Facilitate Medical Innovation (GRIFIN) in Platform Program for Promotion of Genome Medicine (P3GM) of AMED.

## Author contributions

J.H., T.Kato, and N.S. designed the study, performed statistical analysis, and wrote the manuscript. J.H., K.S., T.Kato, and N.S. and contributed to data acquisition. Y.O., M.H., N.S., T.Y., and T.Kadowaki supervised the study. All authors contributed to and approved the final version of the manuscript.

## Competing interests

The authors declare no competing interests.

Supplementary information is available at Human Genome Variation’s website.

