## Supplementary material for "Structural basis of ethnic-specific variants of PAX4 associated with type 2 diabetes"

### Supplementary Note

#### Building of structural models of two PAX4 mutants (p.Arg192Ser and p.Arg192His)

A 3-dimensional structural model of the human PAX4 homeodomain (amino acid residues 162-220) was constructed by homology modeling with SWISS-MODEL[11], using the X-ray crystal structure of human PAX3 (UniProt: P23760) (PDB ID: 3CMY)[12] as a template. Amino acid identity was 50.85% between the PAX4 and PAX3 homeodomains. When the quality of the structural model was evaluated by using QMEAN, the QMEAN Z-score of the PAX4 model was -0.55, indicating that the model was reliable[11]. A structural model of the PAX4 homeodomain binding with DNA was built from PAX3 complexed with DNA (PDB ID: 3CMY)[12] by superimposing PAX4 onto PAX3. Superimposition and merging of coordinate data were performed with Waals (Altif Laboratories, Inc., Tokyo, Japan). Structural models of two PAX4 mutants (p.Arg192Ser and p.Arg192His) were built by using the mutation and energy minimization command of Swiss-Pdb Viewer[13]. Comparison between the structural models of wild-type PAX4 and each mutant was performed with Waals. Coordinate data for the X-ray crystal structures of the human PAX3 (PDB ID: 3CMY)[12] was obtained from the Protein Data Bank: PDB (https://www.rcsb.org).

#### Analysis of structural modifications in the PAX4 mutants (p.Arg192Ser and p.Arg192His)

T2D GWAS meta-analysis recently performed in a Japanese population showed that a previously unreported missense variant of *PAX4* (p.Arg192Ser) reached genome-wide significance[4]. Another previously established independent T2D variant of *PAX4* (p.Arg192His)[14] located at the same amino acid was reported to be associated with reduced C-peptide levels in a genetic study of T2D in Korea[15]. It was reported that an *in vitro* study of the PAX4 p.Arg192His mutant demonstrated impaired repression of the transcription of target genes involved in maintenance of β-cell function compared with wild-type PAX4[16]. To investigate the influence of these *PAX4* mutations, we constructed 3-dimensional structural models of the wild-type human PAX4 homeodomain and the two mutants (Arg192Ser and Arg192His). Then we compared the model of each mutant with that of wild-type PAX4 to identify structural defects. The homeodomain of PAX4 is a conserved DNA-binding module that consists of three α-helices (helix-1, helix-2 and helix-3), and helix-3 of the homeodomain fits into the major groove of DNA (**Fig. 1**). Arg192 is located on the helix-2 and the side chain of Arg192 forms salt-bridges with Glu203 on the helix-3. The residues Arg192 and Glu203 are conserved in the homeodomain family and are considered to contribute to stability of the three helices in the homeodomain structure through the formation of salt bridge[12,17,18]. In addition, Arg192 of PAX4 is thought to bind with the phosphate backbone of double-stranded DNA (**Fig. 1**). The Arg residue corresponding to Arg192 of PAX4 is considered to be involved with DNA binding in other members of the homeodomain family[12,18]. Substitution of Arg192 by Ser or His disrupts the formation of the salt-bridges with Glu203 due to loss of the positively charged arginine side chain (**Fig. 2**), reducing the stability of the homeodomain. In addition, substitution of Arg192 by Ser or His is predicted to disrupt the direct binding with the phosphate backbone of double-stranded DNA (**Fig. 2**). In models of the two mutants (p.Arg192Ser and p.Arg192His) binding to DNA, the distance between the side chain of Ser or His and the phosphate backbone of DNA was 6.57Å and 6.15Å, respectively, indicating these residues could not interact with the phosphate backbone of DNA. Accordingly, the p.Arg192Ser and p.Arg192His variants of PAX4 are deduced to show decreased stability of the homeodomain containing the helices-2 and -3 involved with DNA binding, as well as loss of the ability to bind directly to the phosphate backbone of double-stranded DNA. Thus, these mutations are deduced to affect structural stability of the PAX4 homeodomain and its binding to DNA. It has been reported that missense mutations of the PAX3 homeodomain associated with Waardenburg syndrome are predicted to destabilize the homeodomain or affect DNA binding[12], in agreement with the structural influence of these two PAX4 variants.

**Supplementary Table 1. Variants of genes involved in the MODY pathways associated with T2D in the Japanese population.**

| | | | | | | Japanese | | | | European § | | | | MAF in 1000 Genomes Phase 3 | | |
|---|---|---|---|---|---|---|---|---|---|---|---|---|---|---|---|---|
| LOCUS* | rsID † | CHR | POS | RA | OA | $MAF_{JPN}$ | $P_{JPN}$ | $OR_{JPN}$ | 95%CI | $MAF_{EUR}$ | $P_{EUR}$ | $OR_{EUR}$ | 95%CI | EAS | JPT | EUR |
| *AGPAT9/NKX6-1* | rs201597274 | 4 | 85,301,870 | T | C | 0.044 | 2.1E-12 | 1.17 | 1.12-1.23 | ND | ND | ND | ND | 0.020 | 0.034 | 0.0070 |
| *GCK* | rs2908279 | 7 | 44,174,857 | G | T | 0.40 | 5.3E-12 | 1.07 | 1.05-1.09 | 0.50 | 1.1E-07 | 1.03 | 1.02-1.05 | 0.37 | 0.40 | 0.49 |
| *PAX4* | rs2233580 | 7 | 127,253,550 | T | C | 0.092 | 4.1E-74 | 1.32 | 1.28-1.36 | 0.00050 | 2.9E-01 | 1.62 | 0.66-3.97 | 0.098 | 0.11 | 0 |
| *MNX1* | rs1182389 | 7 | 157,038,803 | G | A | 0.46 | 1.1E-08 | 1.05 | 1.03-1.07 | 0.36 | 1.7E-11 | 1.05 | 1.03-1.06 | 0.47 | 0.47 | 0.37 |
| *HHEX/IDE* | rs12219514 | 10 | 94,466,439 | A | G | 0.13 | 4.8E-39 | 1.18 | 1.15-1.21 | 0.44 | 4.6E-61 | 1.12 | 1.10-1.13 | 0.13 | 0.13 | 0.44 |
| *INS-IGF2_KCNQ1* | rs2237897 ‡ | 11 | 2,858,546 | C | T | 0.38 | 2.6E-168 | 1.30 | 1.27-1.32 | 0.046 | 1.8E-31 | 1.21 | 1.17-1.25 | 0.35 | 0.39 | 0.047 |
| *HNF1A* | rs187150787 | 12 | 121,327,809 | A | G | 0.037 | 6.0E-12 | 1.20 | 1.14-1.27 | ND | ND | ND | ND | 0.020 | 0.024 | 0 |
| *HNF1B* | rs11651052 | 17 | 36,102,381 | A | G | 0.32 | 2.2E-34 | 1.12 | 1.10-1.14 | 0.47 | 8.9E-29 | 1.07 | 1.06-1.09 | 0.25 | 0.33 | 0.47 |
| *HNF4A* | rs16988991 | 20 | 42,989,777 | A | G | 0.45 | 3.8E-09 | 1.05 | 1.04-1.07 | 0.16 | 5.8E-09 | 1.05 | 1.03-1.07 | 0.42 | 0.40 | 0.16 |

CHR, chromosome; POS, position in Human Genome version 19 (hg19), build 37; RA, risk allele; OR, other allele; MAF, minor allele frequency; OR, odds ratio; CI, confidence interval.

* Based on locus information reported by Suzuki et al. (*Nat. Genet.* 2019)[4]

† Variants at primary signals[4] without adjustment for BMI are shown. Information on the same variants from GWAS meta-analysis in a European population[3] without adjustment for BMI is also shown.

‡ The variant rs2237897 is located within 1 Mb of the lead independent variant rs4929965 at the *INS/IGF2* locus in Europeans[3], which is also associated with T2D in Japanese[4] (see Supplementary Table 2).

§ Based on summary statistics reported by Mahajan et al. (*Nat. Genet.* 2018)[3]

**Supplementary Table 2. Variants of genes involved in the MODY pathway associated with T2D in Europeans.**

| | | | | | | European | | | | Japanese ‡ | | | | MAF in 1000 Genomes Phase 3 | | |
|---|---|---|---|---|---|---|---|---|---|---|---|---|---|---|---|---|
| **LOCUS*** | **rsID †** | **CHR** | **POS** | **RA** | **OA** | **$MAF_{EUR}$** | **$P_{EUR}$** | **$OR_{EUR}$** | **95%CI** | **$MAF_{JPN}$** | **$P_{JPN}$** | **$OR_{JPN}$** | **95%CI** | **EAS** | **JPT** | **EUR** |
| *SLC2A2* | rs9873618 | 3 | 170,733,076 | G | A | 0.29 | 4.8E-21 | 1.07 | 1.05-1.08 | 0.20 | 1.0E-02 | 1.03 | 1.01-1.05 | 0.21 | 0.25 | 0.29 |
| *GCK* | rs878521 | 7 | 44,255,643 | A | G | 0.25 | 1.9E-13 | 1.06 | 1.04-1.07 | 0.41 | 2.1E-02 | 1.02 | 1.00-1.04 | 0.35 | 0.36 | 0.25 |
| *MNX1* | rs6459733 | 7 | 156,930,550 | G | C | 0.33 | 2.4E-17 | 1.06 | 1.05-1.07 | 0.46 | 2.7E-07 | 1.05 | 1.03-1.07 | 0.45 | 0.44 | 0.36 |
| *NEUROG3* | rs2642588 | 10 | 71,466,578 | G | T | 0.30 | 2.2E-14 | 1.05 | 1.04-1.07 | 0.082 | 1.9E-01 | 1.02 | 0.99-1.06 | 0.13 | 0.077 | 0.30 |
| *HHEX/IDE* | rs10882101 | 10 | 94,462,427 | T | C | 0.41 | 1.4E-08 | 1.06 | 1.04-1.08 | 0.29 | 1.2E-35 | 1.13 | 1.10-1.15 | 0.28 | 0.33 | 0.43 |
| *INS/IGF2* | rs4929965 | 11 | 2,197,286 | A | G | 0.38 | 4.0E-26 | 1.07 | 1.06-1.09 | 0.081 | 3.1E-13 | 1.12 | 1.09-1.16 | 0.087 | 0.067 | 0.36 |
| *HNF1A* | rs56348580 | 12 | 121,432,117 | G | C | 0.31 | 2.3E-13 | 1.05 | 1.04-1.07 | ND | ND | ND | ND | 0.0050 | 0 | 0.30 |
| *ONECUT1* | rs2456530 | 15 | 53,091,553 | T | C | 0.13 | 5.4E-09 | 1.06 | 1.04-1.08 | 0.49 | 1.3E-02 | 1.02 | 1.00-1.04 | 0.50 | 0.46 | 0.11 |
| *HNF1B* | rs10908278 | 17 | 36,099,952 | T | A | 0.48 | 6.4E-36 | 1.08 | 1.07-1.10 | 0.31 | 6.7E-32 | 1.13 | 1.10-1.15 | 0.23 | 0.32 | 0.48 |
| *NKX2.2* | rs13041756 | 20 | 21,466,795 | C | T | 0.11 | 1.4E-08 | 1.06 | 1.04-1.08 | 0.20 | 8.8E-01 | 1.00 | 0.98-1.02 | 0.21 | 0.23 | 0.10 |
| *HNF4A* | rs1800961 | 20 | 43,042,364 | T | C | 0.035 | 2.3E-22 | 1.18 | 1.15-1.23 | 0.010 | 7.7E-05 | 1.18 | 1.09-1.29 | 0.016 | 0.0048 | 0.037 |

CHR, chromosome; POS, position in Human Genome version 19 (hg19), build 37; RA, risk allele; OR, other allele; MAF, minor allele frequency; OR, odds ratio; CI, confidence interval.

* Based on locus information reported by Mahajan et al. (*Nat. Genet.* 2018)[3]

† Variants at primary signals[3] without adjustment for BMI are shown. Information on the same variants from GWAS meta-analysis in a Japanese population[4] without adjustment for BMI is also shown.

‡ Based on summary statistics reported by Suzuki et al. (*Nat. Genet.* 2019)[4]
